# Asperosaponin VI inhibits LPS-induced inflammatory response by activating PPAR-γ pathway in primary microglia

**DOI:** 10.1101/2020.03.15.992453

**Authors:** Jinqiang Zhang, Saini Yi, Chenghong Xiao, Yahui Li, Chan Liu, Weike Jiang, Changgui Yang, Tao Zhou

**Affiliations:** Guizhou University of Traditional Chinese Medicine, Guiyang 550025, China

**Keywords:** Asperosaponin VI, Microglia, Neuroinflammation, lipopolysaccharide

## Abstract

Microglia cells are the main mediators of neuroinflammation. Activation of microglia often aggravates the pathological process of various neurological diseases. Natural chemicals have unique advantages in inhibiting microglia-mediated neuroinflammation and improving neuronal function. Here, we examined the effects of asperosaponin VI (ASA VI) on LPS-activated primary microglia. Microglia were isolated from mice and pretreated with different doses of ASA VI, following lipopolysaccharide (LPS) administration. Activation and inflammatory response of microglia cells were evaluated by q-PCR, immunohistochemistry and ELISA. Signaling pathways were detected by western blotting. We found that the ASA VI inhibited the morphological expansion of microglia cells, decreased the expression and release of proinflammatory cytokines, and promoted the expression of antiinflammatory cytokines in a dose-dependent manner. ASA VI also activated PPAR-γ signaling pathway in LPS-treated microglia. The anti-inflammatory effects of ASA VI in microglia were blocked by treating PPAR-γ antagonist (GW9662). These results showed that ASA VI promote the transition of microglia cells from proinflammatory to anti-inflammatory by regulating PPAR-γ pathway.

## 1. Introduction

Microglia are the only glial cells that derive from a restricted subpopulation of yolk sac erythromyeloid progenitors. Microglia play central roles in immune surveillance and inflammatory-related neuropathology (Yirmiya et al., 2015). Dysfunction of microglia has been associated with a variety of psychiatric disorders, including depression, autism, and schizophrenia (Yirmiya et al., 2015, Brites and Fernandes,2015). The cascade of microglial activation could promote the synthesis and secretion of a large number of inflammatory mediators, resulting in nerve damage and accelerating the pathological progression of neurodegenerative diseases (Zhang et al., 2016, Zhang et al., 2018b). Inhibiting the microglia-meiated neuroinflammation will be an important strategy for Alzheimer’s disease and depression (Zhang et al.,2018b).

Peroxisome proliferator activated receptor gamma (PPAR-γ) plays an important role in regulating transcription expression of anti-inflammatory cytokines (Han et al., 2017). Studies have shown that PPAR-γ agonists (pioglitazone or rosiglitazone) can switch the transition of microglia cells from a proinflammatory to an anti-inflammatory state (Zhao et al., 2016, Wen et al., 2018). Natural compounds have broad prospects in neuroimmune regulation. Many studies have shown that natural products, including ginsenoside Rb1 and salvianolic acid B, etc. have a good inhibitory effect on microglia-mediated neuroinflammation, which could promote the expression of anti-inflammatory cytokines by activating PPAR-γ signaling pathways (Lu et al., 2017, Zhang et al.,2018a).

Dipsaci Radix (DR), the desiccative radix of *Dipsacus asper*., is an important and common traditional Chinese medicine, has a long application history on clinical. The current research showed DR has multiple pharmaco-activities, and the main effective part is the saponins (Gao et al., 2016). Among them, one of the highest content of saponins is asperosaponin VI (ASA VI) (Ding et al., 2019). The effect of ASA VI on proliferation and differentiation of osteoblasts has been widely reported (Gao et al., 2016, Liu et al., 2019). However, the role of ASA VI in innate immune regulation is unclear. In this study, we identified the regulatory effect of ASA VI on microglia cells and explored its molecular mechanism, laying a foundation for further studies on development of its pharmacological potential.

## 2. Materials and methods

### 2.1 Primary microglia culture

Microglia cells were cultured as previously described (Zhang et al.,2017). In brief, mixed glial cells were isolated from the brain of P0-P3 C57BL/6J mice. These mixed glial cells were maintained for 10–14 d in DMEM (Invitrogen Gibco, USA) containing 10% FBS (Invitrogen Gibco, USA). Microglia cells are separated from the mixed glia by shock.

### 2.2 LPS administration and pharmacological intervention

The asperosaponin VI (ASA VI) standard (purity = 99.92%) was purchased from Chengdu Alfa Biotechnology Co., Ltd. The asperosaponin VI was dissolved in sterile phosphate belanced solution (PBS). The microglia were pretreated with ASA VI of 10 μM, 50 μM, 100 μM and 200 μM. After 30 min, these microglia were treated 100 ng/mL LPS (Sigma, USA) for 24 h. Following the immunocytochemistry, RT-PCR analysis, ELISA and western blot analysis were performed.

### 2.3 Immunocytochemistry and image analysis

The purified microglia cells were cultured in 24 well plates at 1 × 10^5^ cells. These microglia cells were incubated with primary antibodies (mouse anti-Iba1 antibody, abcam, 1:400; rabbit anti-CD68 antibody, AbD Serotec, 1:100; rabbit anti-iNOS antibody, abcam,1:50; rabbit anti-CD206 antibody, abcam, 1:200; rabbit anti-PPAR-γ antibody, Cell Signaling Technology, 1:200) for 24 h and the secondary antibodies (anti-rabbit IgG-conjugated Alexa Fluorochrome or anti-mouse IgG conjugated Alexa Fluorochrome, Invitrogen; 1:500) for 2 h at room temperature.

Fluorescence micrographs of microglia cells were captured by a fluorescence microscopy (Olympus BX51). Image J software (version 1.45 J) was used to measure the area, perimeter and fluorescence intensity of microglia.

### 2.4 Quantitative PCR

The purified microglia were cultured in 6-well plates at 1 × 10^6^ cells. 24 h after LPS treatment, total RNA was extracted using Trizol (Invitrogen Life Technologies, USA). Using the reverse transcription kit (TaKaRa, Japan) to get the cDNA in strict accordance with the steps in the experimental instructions. The RT-PCR reaction mixture contains 1 μL of template cDNA, 5 μL MasterMix and 1 μL primer (Sangon Biotech, Sichuan, China), add DEPC water to a total reaction volume of 10 μL. After mixing, put it into the 7500 Real-Time PCR System (Applied Biosystems, USA). The internal reference gene is β-actin and the expression of related genes are calculated according to the method of -ΔΔ Ct. Primer sequences have been listed in Table S1.

### 2.5 Enzyme-linked immunosorbent assay (ELISA)

The purified microglia cells were cultured in 6-well plates at 1 × 10^6^ cells. The supernatant is collected and centrifuged to remove the precipitate for testing secretion of inflammatory cytokines. BCA (Sangon Biotech, Sichuan, China) was used to determine the concentration of total protein, diluted to the same concentration, and then operated in strict accordance with the instructions of the ELISA Kit (BOSTER, Wuhan, China) to determine the levels of IL-10, IL-1β and TNF-α.

### 2.6 Western blot analysis

The purified microglia cells were cultured at 1 × 10^6^ cells and lysed using RIAP lysis (Solarbio, China). The protein concentration of the lysates solution was tested by BCA method. 12% SDS polyacrylamide gel was used to resolve protein. The PVDF membranes which transformed with protein were blocked with 5% skimmed milk. Primary antibodies (rabbit anti-PPAR-γ antibody, Cell Signaling Technology, 1:1000) and the secondary antibodies (goat anti-rabbit IgG) were incubated with the PVDF membranes. The proteins levels of PVDF membranes were measured with the chemiluminescence detection system (Amersham, Berkshire, UK).

### 2.7 Statistic analysis

All of the data are showed as the Means ±SEM. Graph drawing was used GraphPad Prism (version 7.0). SPSS software was used for significance analysis. Potential differences between the mean values were evaluated using Independent-Sample t test and one/two-way analysis of variance (ANOVA) followed by Tukey’s multiple comparison test for post hoc comparisons. P<0.05 was considered to indicate a statistically significant difference. Each sample was repeated 3 times for q-PCR, ELISA and western blot, 5 immunofluorescence images of each simple were used to image analysis. The mean value of the parallel repeated data was used for significance statistical analysis.

## 3. Results

### 3.1 ASA VI inhibits LPS-induced activation of microglia and proinflammatory production

The microglia were treated with LPS to induce the inflammatory response. The results from immunohistochemistry showed that LPS treatment results in the morphological change of microglia, including the increased area, perimeter, TI value and CD68^+^ area. The ASA VI inhibited LPS-induced activation of microglia in a dose-dependent manner. When the microglia were pretreated with ASA VI (100 μM, 200 μM) before LPS administration, the area, perimeter, TI value and CD68^+^area of microglia were significantly inhibited. The 10 μM and 50 μM ASA VI were not significantly affect the morphology of microglia (Fig. 1A-1F).

**Fig. 1.**
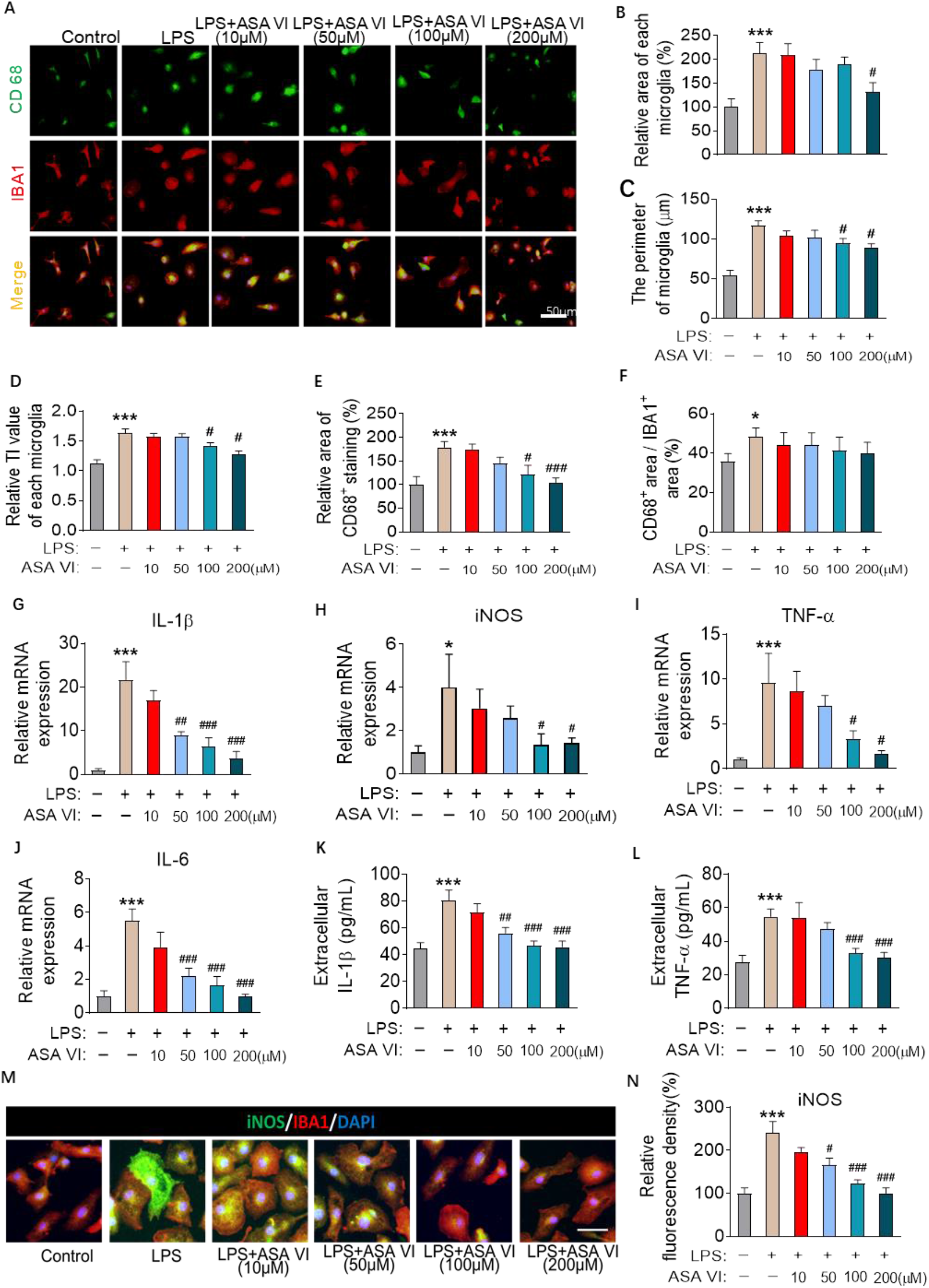
ASA VI inhibits LPS-induced activation of microglia and proinflammatory production. **A**: Immunohistochemistry detects the morphological changes and phagolysosome of microglia cells in LPS-induced primary microglia after pretreating different doses of ASA VI. Microglia cells are labeled with IBA1 (red), phagolysosome is labeled with CD68 (green), and nucleus is labeled DAPI. Scale bar, 50μm. **B**: Quantification of the relative area of each microglia. Data are standardized to control group. **C**: Quantification of the perimeter of each microglia. **D**: Quantification of the TI value of each microglia. **E**: Quantification of the relative area of CD68^+^ staining in each microglia. Data are standardized to control group. **F**: Quantification of the percentage of CD68^+^ staining out of IBA1^+^ staining in each microglia. **G-J**: Quantitative PCR detects the mRNA expression of proinflammatory cytokines (IL-1β, iNOS, TNF-α and IL-6). Data are showed the fold change relative to control group. **K** and **L**: ELISA detects the extracellular protein levels of proinflammatory cytokines (IL-1β and TNF-α). **M**: Immunohistochemistry detects iNOS (green) expression in LPS-induced primary microglia after pretreating different doses of ASA VI. Scale bar, 10μm. **N**: Quantification of the relative fluorescence intensity of iNOS. Data are standardized to control group. Data are mean ±SEM (n =3-5 per group), each sample was repeated 3 times for q-PCR and ELISA, 5 immunofluorescence images of each simple were used to analysis. * P < 0.05, ** P < 0.01, *** P < 0.005 when compared with control group, ^#^ P < 0.05, ^##^ P < 0.01, ^###^ P < 0.005 when compared with LPS group (one-way ANOVA with LSD test).

The expression of inflammatory cytokines is usually synchronized with changes in microglial morphology. The pro-inflammatory cytokines expressions were evaluated in LPS-treated microglia. The results from q-PCR showed that the 100 μM and 200 μM ASA VI significantly suppressed the the gene expression of IL-1β, iNOS, TNF-α and IL-6 in microglia. And the 50 μM ASA VI only suppressed the expression of IL-1β and IL-6 but not iNOS and TNF-α (Fig. 1G-1J). ELISA was performed to measure the concentration of IL-1β and TNF-α which secreted by microglia cells in the medium. The results showed that 100 μM and 200 μM ASA VI significantly reduced the secretion of IL-1β and TNF-α in LPS-treated microglia (Fig. 1K and 1L). The immunohistochemistry was performed to measure the intracellular expression of iNOS in microglia. The result showed 50 μM, 100 μM and 200 μM ASA VI significantly inhibited the intracellular expression of iNOS (Fig. 1M and 1N).

### 3.2 ASA VI promotes anti-inflammatory production in LPS-induced microglia

We next examined the effects of ASA VI on antiinflammatory cytokines expression in LPS-induced microglia. The results from q-PCR showed that the 100 μM and 200 μM ASA VI significantly increased the CD206 and IL-10 expression in LPS-induced microglia. The 50 μM ASA VI upregulated the expression of IL-10 but not CD206 (Fig. 2A and 2B). ELISA was performed to measure the concentration of IL-10 which secreted by microglia cells in the medium. The results showed that 100 μM ASA VI significantly upregulated the secretion of IL-10 in LPS-treated microglia (Fig. 2C). The result from immunohistochemistry showed 50 μM, 100 μM and 200 μM ASA VI significantly increased the relative fluorescence intensity of CD206 in LPS-treated microglia (Fig. 2D and 2E).

**Fig. 2.**
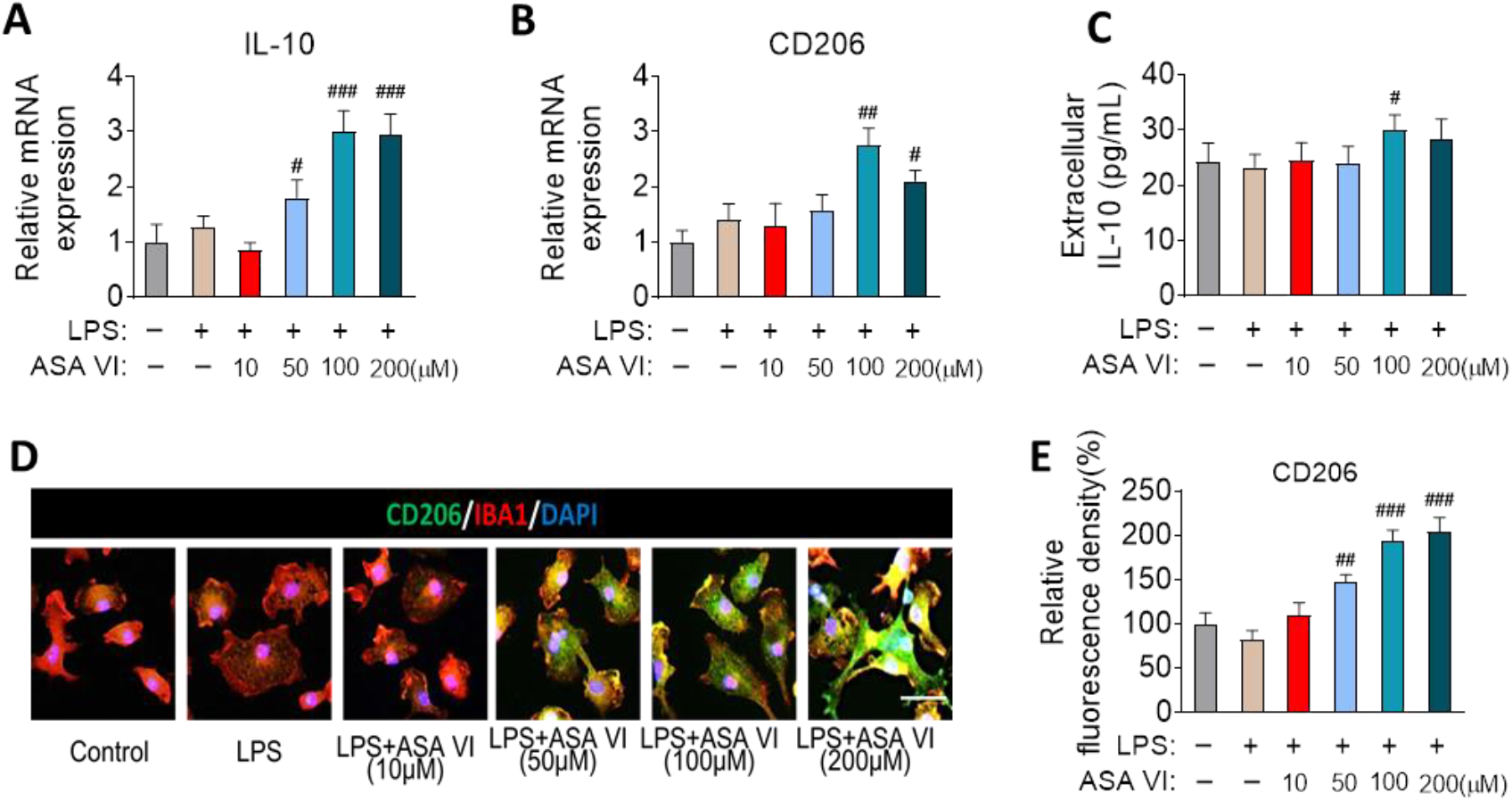
ASA VI promotes anti-inflammatory production in LPS-induced microglia. **A** and **B**: Quantitative PCR detects the mRNA expression of anti-inflammatory cytokines (IL-10 and CD206). Data are showed the fold change relative to control group. **C**: ELISA detects the extracellular protein levels of anti-inflammatory cytokines (IL-10). **D**: Immunohistochemistry detects CD206 (green) expression in LPS-induced primary microglia after pretreating different doses of ASA VI. Scale bar, 10μm. **E**: Quantification of the relative fluorescence intensity of CD206. Data are standardized to control group. Data are mean ± SEM (n =3-5 per group), each sample was repeated 3 times for q-PCR and ELISA, 5 immunofluorescence images of each simple were used to analysis. ^#^ P < 0.05, ^##^ P < 0.01, ^###^ P < 0.005 when compared with LPS group (one-way ANOVA with LSD test).

### 3.3 ASA VI increases the expression and nuclear translocation of PPAR-γ in a dose manner

To further explore the molecular mechanism by which ASA VI inhibit microglial activation, we examined the transcriptional expression and activation of nuclear transcription factors PPAR-γ in ASA VI-treated microglia. We found that ASA VI increased the transcriptional expression of PPAR-γ in LPS-induced microglia (Fig. 3A). And the IL-1β expression was negatively correlated with the PPAR-γ transcriptional expression (Fig. 3B). We next examined nuclear translocation of PPAR-γ after ASA VI treatment in LPS-induced microglia. The results showed ASA VI significantly increased the percentage of nuclear translocation of PPAR-γ in LPS-induced microglia (Fig. 3C and 3D).

**Fig. 3.**
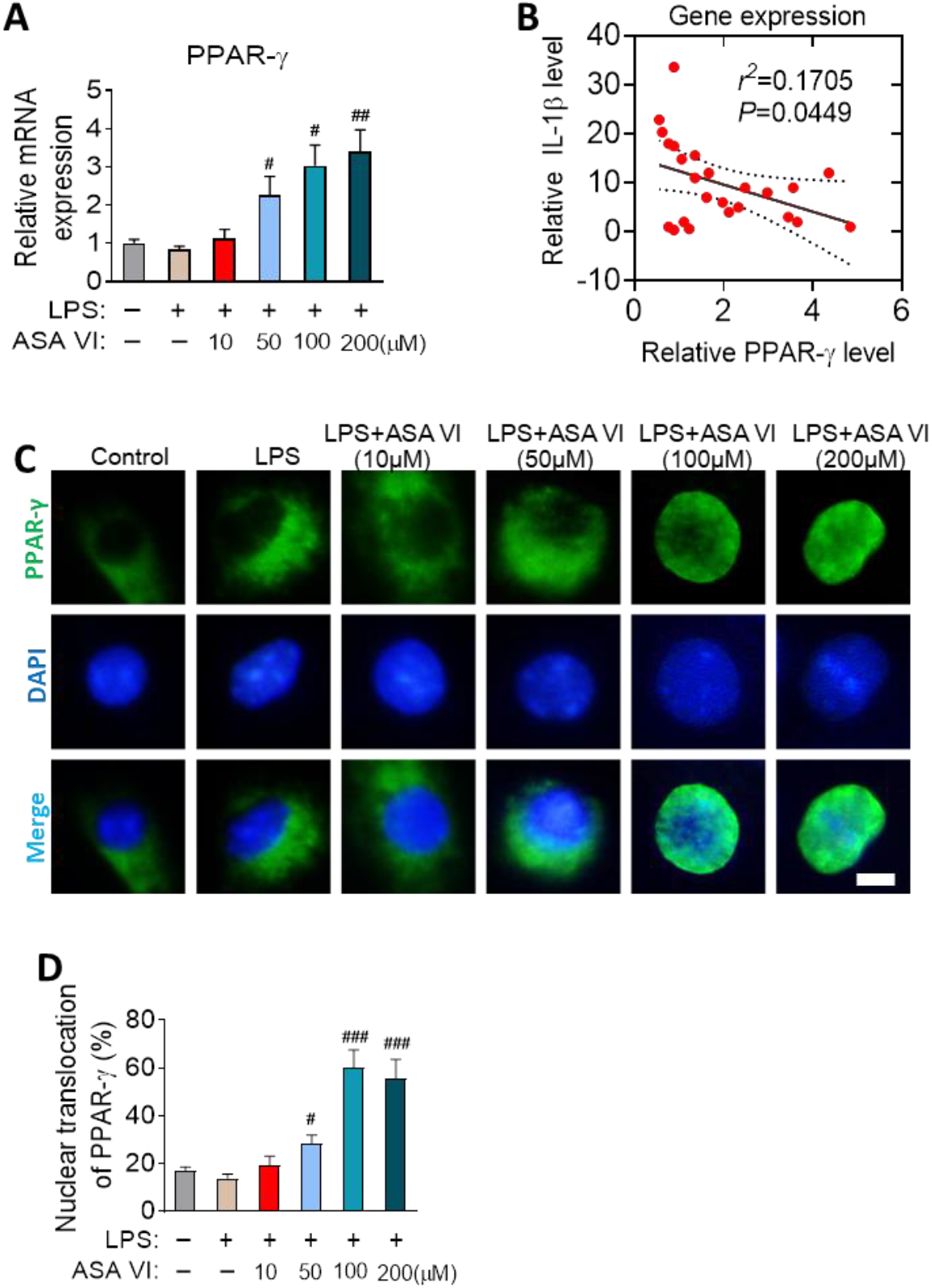
ASA VI increases the expression and nuclear translocation of PPAR-γ in a dose manner. **A**: Quantitative PCR detects the mRNA expression of PPAR-γ. Data are showed the fold change relative to control group. **B**: Correlation analysis between the gene expression of IL-1β and PPAR-γ. 24 simples that from 6 groups (4 simple each group) were used to analysis. Each dot represents a sample. *r^2^*=0.1705, *P*=0.0449. **C**: Immunohistochemistry detects the nuclear (blue) translocation of PPAR-γ (green) in LPS-induced primary microglia after pretreating different doses of ASA VI. Scale bar, 2μm. **D**: Quantification of the percentage of nuclear translocation of PPAR-γ. The ratio nuclear translocation of PPAR-γ was evaluated by the PPAR-γ^+^-DAPI^+^ area out of DAPI^+^ area. Data are mean ± SEM (n =3-5 per group), each sample was repeated 3 times for q-PCR, 5 immunofluorescence images of each simple were used to analysis. *** P < 0.005 when compared with control group, ^#^ P < 0.05, ^##^ P < 0.01, ^###^ P < 0.005 when compared with LPS group (one-way ANOVA with LSD test).

### 3.4 The anti-inflammatory effects of ASA VI in microglia can be blocked by GW9662

In order to further verify that nuclear transcription factors PPAR-γ are necessary for the anti-inflammatory activity of ASA VI in microglia, the antagonists of PPAR-γ (GW9662) are used to block the activation of PPAR-γ signaling pathways in ASA VI-treated microglia (Fig. 4A-4E). The results showed that the effect of ASA VI on microglial morphology was blocked by GW9662 treatment (Fig. 4F and 4G). GW9662 treatment also blocked the inhibitory effect of ASA VI on proinflammatory cytokines (IL-1β) expression and secretion in LPS-induced microglia (Fig. 4H and 4I). Moreover, the increase in expression and secretion of IL-10 induced by ASA VI were blocked by GW9662 treatment in LPS-treated microglia (Fig. 4J and 4K).

**Fig. 4.**
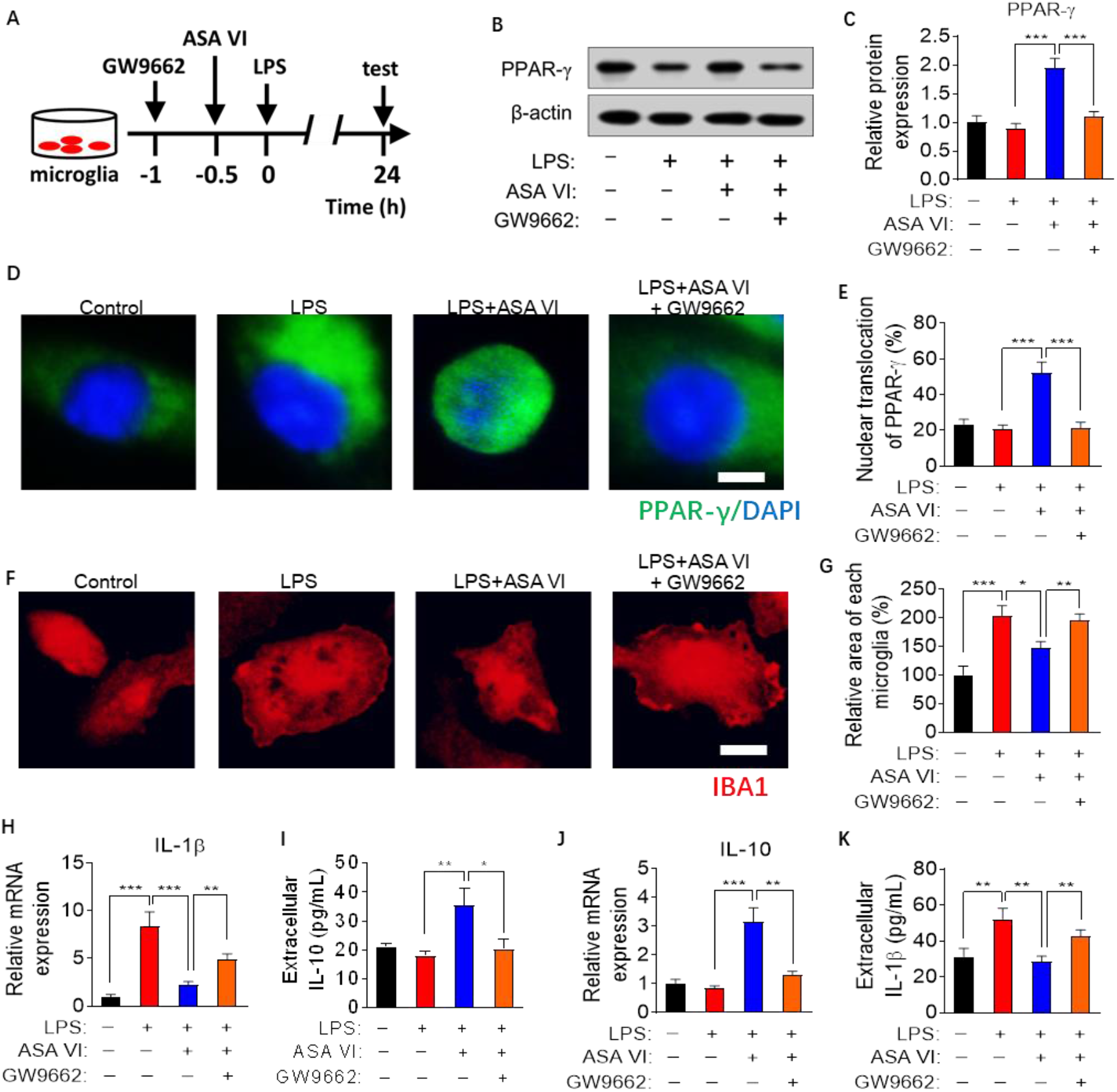
The anti-inflammatory effects of ASA VI were blocked by GW9662. **A**: Schematic diagram for blocking PPAR-γ signaling pathway with GW9662 in ASA VI-treated microglia cells. **B**: Western blotting detects PPAR-γ when pretreated GW9662 before treat ASA VI (40 μM) in LPS-induced primary microglia. **C**: Quantification of the relative protein level of PPAR-γ when pretreated GW9662 before treat ASA VI in LPS-induced primary microglia. PPAR-γ was normalized β-actin. **D**: Immunohistochemistry detects the nuclear (blue) translocation of PPAR-γ (green) when pretreated GW9662 before treat ASA VI in LPS-induced primary microglia. Scale bar, 2μm. **E**: Quantification the percentage of nuclear translocation of PPAR-γ. The ratio nuclear translocation of PPAR-γ was evaluated by the PPAR-γ^+^-DAPI^+^ area out of DAPI^+^ area. **F**: Immunohistochemistry detects the morphological changes of microglia cells when pretreated GW9662 before treat ASA VI in LPS-induced primary microglia. Microglia cells are labeled with IBA1 (red). Scale bar, 10μm. **G**: Quantification of the relative area of microglia when pretreated GW9662 before treat ASA VI in LPS-induced primary microglia. **H:** Quantitative PCR detects the mRNA expression of proinflammatory cytokines (IL-1β). Data are showed the fold change relative to control group. **I:** ELISA detects the extracellular protein levels of proinflammatory cytokines (IL-1β). **J**: Quantitative PCR detects the mRNA expression of anti-inflammatory cytokines (IL-1β). Data are showed the fold change relative to control group. **K**: ELISA detects the extracellular protein levels of anti-inflammatory cytokines (IL-10). Data are mean ±SEM (n =3-5 per group), each sample was repeated 3 times for western blotting, 5 immunofluorescence images of each simple were used to analysis. * P < 0.05, ** P < 0.01, *** P < 0.0 5 (two-way ANOVA with LSD test).

## 4. Discussion

Although there has been no report on the neuroimmunomodulation of ASA VI, according to the reducing swelling, antioxidant and relieving pain properties of Dipsaci Radix, we boldly speculated ASA VI that the chemical components with the highest content in *Dipsaci Radix* might have the effect of inhibiting inflammatory response. And for all we know this study is the first to demonstrate ASA VI suppresses LPS-induced activation of microglia by activating PPAR-γ pathway.

Microglia are thought to be macrophages that resident in the brain (Wolf et al., 2017). They are a class of innate immune cells which play an important role in neuroimmunomodulation (Colonna and Butovsky, 2017). In recent years, the role of microglia cells in neurological diseases has received more and more attention. Inhibiting the activation of microglia cells is one of the effective ways to prevent neuroinflammatory injury (Zhang et al., 2016). Activation of microglia cells is usually characterized by increased cell body and phagolysosome, as well as increased release of proinflammatory cytokines (Zhang et al., 2017). In this study, LPS-treated microglia showed significant activation characteristics. The ASA VI suppressed the LPS-induced increase in the diameter and area of microglia, area of phagolysosome, and synthesis and secretion of pro-inflammatory mediators. These data indicated that ASA VI inhibits LPS-induced activation of microglia and proinflammatory production.

In addition to inhibiting proinflammatory cytokines, the upregulating the anti-inflammatory cytokines expression also plays an important role in anti-inflammatory effect of drugs (Schain and Kreisl, 2017, Carlessi et al., 2019). In this study, we found that ASA VI could increase the expression and secretion of IL-10, a classic anti-inflammatory cytokine. The anti-inflammatory phenotypic marker (CD206) of microglia cells also increased significantly after ASA VI treatment. These findings suggested that ASA VI promoted the phenotypic transition of LPS-treated microglia from M1 to M2.

PPAR-γ, as one of nuclear transcription factors, usually needs to bind to their corresponding ligands and transfer to the nucleus to initiate transcription (Yao et al., 2019). Activation of PPAR-γ usually inhibits pro-inflammatory signaling pathways, such as TLR4/NF-κb, JAK/STAT1, etc., thereby inhibiting the gene transcriptional expression of pro-inflammatory cytokines (Machado et al., 2019). On the other hand, activation of PPAR-γ increases the gene transcriptional expression of anti-inflammatory genes (Lecca et al., 2018, Tian et al., 2019). In this study, we found ASA VI promoted the transcriptional expression of PPAR-γ gene in LPS-treated microglia. Simultaneously, ASA VI promotes nuclear transfer of PPAR-γ in LPS-induced microglia. These data indicated that ASA VI activated the PPAR-γ signaling pathway in LPS-induced microglia. Activation of PPAR-γ negatively regulated the synthesis and secretion of proinflammatory cytokins.

Finally, we used the antagonists of PPAR-γ (GW9662) to block activation of PPAR-γ signaling pathways in ASA VI-treated microglia. Our data showed that GW9662 blocked inhibitory effect of ASA VI on microglia activation and proinflammatory cytokines expression and secretion, as well as decreased anti-proinflammatory cytokines expression in LPS-induced microglia. These results indicated that activation of PPAR-γ is necessary for anti-inflammatory activity of ASA VI, which plays an important role in ASA VI-promoted phenotype from M1 to M2.

In conclusion, the ASA VI promotes the transformation of LPS-induced microglia from proinflammatory to anti-inflammatory phenotype by activating PPAR-γ pathway.

## Acknowledgments

This study was funded by Key project at central government level: The ability establishment of sustainable use for valuable Chinese medicine resources (Grant No. 2060302), Department of Science and Technology of Guizhou High-level Innovative Talents ([2018]5638) and Guizhou science and technology plan project ([2019]5611). Thanks to professor Lanping Guo from Resource Center of Chinese Academy of Traditional Chinese Medicine for her support and help.

---

**Table S1.**
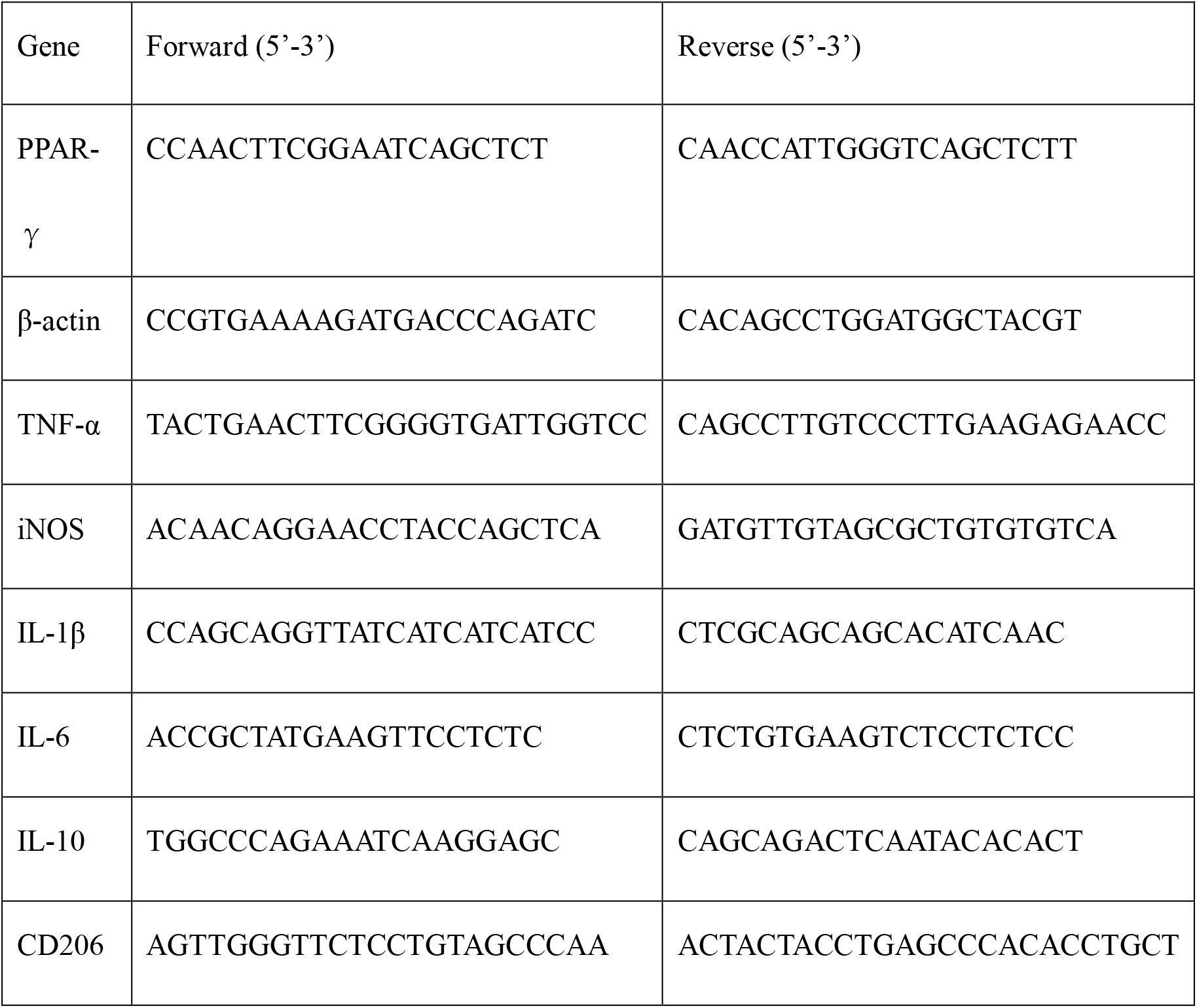
Primers of RT-PCR.

## Reference

Brites, D. & Fernandes, A. 2015. Neuroinflammation and Depression: Microglia Activation, Extracellular Microvesicles and microRNA Dysregulation. Front Cell Neurosci, 9, 476.

Carlessi, A. S., Borba, L. A., Zugno, A. I., Quevedo, J. & Reus, G. Z. 2019. Gut-microbiota-brain axis in depression: The role of neuroinflammation. Eur J Neurosci.

Colonna, M. & Butovsky, O. 2017. Microglia Function in the Central Nervous System During Health and Neurodegeneration. Annu Rev Immunol, 35, 441–468.

Ding, X., Li, W., Chen, D., Zhang, C., Wang, L., Zhang, H., Qin, N. & Sun, Y. 2019. Asperosaponin VI stimulates osteogenic differentiation of rat adipose-derived stem cells. Regen Ther, 11, 17–24.

Gao, J., Zhou, C., Li, Y., Gao, F., Wu, H., Yang, L., Qiu, W., Zhu, L., Du, X., Lin, W., Huang, D., Liu, H., Liang, C. & Luo, S. 2016. Asperosaponin VI promotes progesterone receptor expression in decidual cells via the notch signaling pathway. Fitoterapia, 113, 58–63.

Han, L., Shen, W. J., Bittner, S., Kraemer, F. B. & Azhar, S. 2017. PPARs: regulators of metabolism and as therapeutic targets in cardiovascular disease. Part II: PPAR-beta/delta and PPAR-gamma. Future Cardiol, 13, 279–296.

Lecca, D., Janda, E., Mulas, G., Diana, A., Martino, C., Angius, F., Spolitu, S., Casu, M. A., Simbula, G., Boi, L., Batetta, B., Spiga, S. & Carta, A. R. 2018. Boosting phagocytosis and anti-inflammatory phenotype in microglia mediates neuroprotection by PPARgamma agonist MDG548 in Parkinson’s disease models. 175, 3298–3314.

Liu, K., Liu, Y., Xu, Y., Nandakumar, K. S., Tan, H., He, C., Dang, W., Lin, J. & Zhou, C. 2019. Asperosaponin VI protects against bone destructions in collagen induced arthritis by inhibiting osteoclastogenesis. Phytomedicine, 63, 153006.

Lu, H., Zhou, X., Kwok, H. H., Dong, M., Liu, Z., Poon, P. Y., Luan, X. & Ngok-Shun Wong, R. 2017. Ginsenoside-Rb1-Mediated Anti-angiogenesis via Regulating PEDF and miR-33a through the Activation of PPAR-gamma Pathway. Front Pharmacol, 8, 783.

Machado, M. M. F., Bassani, T. B., Coppola-SEGOVIA, V., Moura, E. L. R., Zanata, S. M., Andreatini, R. & Vital, M. 2019. PPAR-gamma agonist pioglitazone reduces microglial proliferation and NF-kappaB activation in the substantia nigra in the 6-hydroxydopamine model of Parkinson’s disease. Pharmacol Rep, 71, 556–564.

Schain, M. & Kreisl, W. C. 2017. Neuroinflammation in Neurodegenerative Disorders-a Review. Curr Neurol Neurosci Rep, 17, 25.

Tian, Y., Yang, C., Yao, Q., Qian, L., Liu, J., Xie, X., Ma, W., Nie, X., Lai, B., Xiao, L. & Wang, N. 2019. Procyanidin B2 Activates PPARgamma to Induce M2 Polarization in Mouse Macrophages. Front Immunol, 10, 1895.

Wen, L., You, W., Wang, H., Meng, Y., Feng, J. & Yang, X. 2018. Polarization of Microglia to the M2 Phenotype in a Peroxisome Proliferator-Activated Receptor Gamma-Dependent Manner Attenuates Axonal Injury Induced by Traumatic Brain Injury in Mice. J Neurotrauma, 35, 2330–2340.

Wolf, S. A., Boddeke, H. W. & Kettenmann, H. 2017. Microglia in Physiology and Disease. Annu Rev Physiol, 79, 619–643.

Yao, X., Jiang, Q., Ding, W., Yue, P., Wang, J., Zhao, K. & Zhang, H. 2019. Interleukin 4 inhibits high mobility group box-1 protein-mediated NLRP3 inflammasome formation by activating peroxisome proliferator-activated receptor-gamma in astrocytes. Biochem Biophys Res Commun, 509, 624–631.

Yirmiya, R., Rimmerman, N. & Reshef, R. 2015. Depression as a microglial disease. Trends Neurosci, 38, 637–658.

Zhang, J., Xie, X., Tang, M., Zhang, J., Zhang, B., Zhao, Q., Han, Y., Yan, W., Peng, C. & You, Z. 2017. Salvianolic acid B promotes microglial M2-polarization and rescues neurogenesis in stress-exposed mice. Brain Behav Immun, 66, 111–124.

Zhang, J. Q., Wu, X. H., Feng, Y., Xie, X. F., Fan, Y. H., Yan, S., Zhao, Q. Y., Peng, C. & You, Z. L. 2016. Salvianolic acid B ameliorates depressive-like behaviors in chronic mild stress-treated mice: involvement of the neuroinflammatory pathway. Acta Pharmacol Sin, 37, 1141–53.

Zhang, L., Gao, J., Tang, P., Chong, L., Liu, Y., Liu, P., Zhang, X., Chen, L. & Hou, C. 2018a. Nuciferine inhibits LPS-induced inflammatory response in BV2 cells by activating PPAR-gamma. Int Immunopharmacol, 63, 9–13.

Zhang, L., Zhang, J. & You, Z. 2018b. Switching of the Microglial Activation Phenotype Is a Possible Treatment for Depression Disorder. Front Cell Neurosci, 12, 306.

Zhao, Q., Wu, X., Yan, S., Xie, X., Fan, Y., Zhang, J., Peng, C. & You, Z. 2016. The antidepressant-like effects of pioglitazone in a chronic mild stress mouse model are associated with PPARgamma-mediated alteration of microglial activation phenotypes. J Neuroinflammation, 13, 259.

